# Altered Cortical Activation Associated with Mirror Overflow Driven by Non-Dominant Hand Movement in Attention-Deficit/Hyperactivity Disorder

**DOI:** 10.1101/2021.04.23.441107

**Authors:** Yu Luo, Christine Chen, Jack H Adamek, Deana Crocetti, Stewart H Mostofsky, Joshua B Ewen

**Affiliations:** School of Biological Science and Medical Engineering, Beihang University, Beijing, BJ, China; Kennedy Krieger Institute, Baltimore, MD, USA; Department of Neurology, Johns Hopkins University School of Medicine, Baltimore, MD, USA; Department of Psychiatry and Behavioral Sciences, Johns Hopkins University School of Medicine, Baltimore, MD, USA

**Keywords:** Mirror overflow, Event-related desynchronization, Attention-deficit/hyperactivity disorder, Hand skill asymmetry

## Abstract

Mirror overflow is involuntary movement that accompanies unilateral voluntary movement on the opposite side of the body, and is commonly seen in Attention-Deficit/Hyperactivity Disorder (ADHD). Children with ADHD show asymmetry in mirror overflow between dominant and non-dominant hand, yet there are competing mechanistic accounts of why this occurs. Using EEG during a sequential, unimanual finger-tapping task, we found that children with ADHD exhibited significantly more mirror overflow than typically developing (TD) controls, especially during the tapping of the non-dominant hand. Furthermore, source-level EEG oscillation analysis revealed that children with ADHD showed decreased alpha (8-12 Hz) event-related desynchronization (ERD) compared with controls in both hemispheres, but only during tapping of the non-dominant hand. Moreover, only the ERD ipsilateral to the mirror overflow during non-dominant hand movement correlated with both magnitude of overflow movements and higher ADHD symptom severity (Conners ADHD Hyperactivity/Impulsiveness scale) in children with ADHD. TD controls did not show these relationships. Our findings suggest that EEG differences in finger-tapping in ADHD are related primarily to voluntary movement in the non-dominant hand. Our results are also consistent with the Ipsilateral Corticospinal Tract (CST) Hypothesis, which posits that the atypical persistence of mirror overflow in ADHD may originate in the sensorimotor areas ipsilateral to mirror overflow and be transmitted via non-decussating CST fibers.

## 1. INTRODUCTION

Attention-deficit/hyperactivity disorder (ADHD) is the most common and persistent neuropsychiatric condition in childhood, characterized by age-inappropriate hyperactivity, and/or impulsiveness, and by deficits in attention (Lionel et al., 2011). ADHD affects approximately4-9% of school-age children worldwide, incurs high medical costs, increases the risk for academic failure, adult mental illness, substance abuse and criminal behavior (Canino et al., 2004; Swanson et al., 2006). Long-term follow-up studies suggest that even the most aggressive treatment regimens fail to prevent high rates of adverse social outcomes (Polanczyk et al., 2007; Szatmari et al., 1989).

Mirror overflow, defined as the involuntary and unintentional movement of the homologous muscles accompanying voluntary actions of the contralateral body part (Cox et al., 2012), has been commonly reported in ADHD and other neurodevelopmental disorders (e.g., schizophrenia, autism) (MacNeil et al., 2011; McAuliffe et al., 2020; Mostofsky et al., 2003; Mostofsky et al., 2009; Mostofsky et al., 2006). However, the physiological origins of mirror overflow remain controversial. Two competing models form a framework to explain the cortical origins of mirror overflow. The Transcallosal Hypothesis attributes the presence of mirror overflow to dysregulation of transcallosal interhemispheric inhibition and facilitation from the hemisphere contralateral to the overflow, whereas the Ipsilateral Corticospinal Tract (CST) Hypothesis states that the hemisphere ipsilateral to overflow movements is involved in the production of mirror overflow via non-decussating CST fibers (Hoy et al., 2004). Because the motor system is relatively well characterized, compared with the biology underlying more complex behavioral phenomena of ADHD, investigating the brain basis of motor atypicalities could further our understanding the neurobiology of the disorder as a whole and aid in the identification of biomarkers helpful for improving targeted diagnosis and treatment.

Several studies have proposed both empiric and putative neurobiologically mechanistic associations between mirror overflow and the diagnostic behavioral manifestations of ADHD (Mostofsky et al., 2003; Mostofsky et al., 2009; Mostofsky et al., 2006). Specially, excessive motor overflow significantly predicts performance on measures of behavioral response inhibition in children with ADHD such that greater overflow is associated with worse inhibition (Mostofsky et al., 2003) and is correlated with greater Hyperactive/Impulsive (H/I) ADHD symptoms (MacNeil et al., 2011).

More nuanced parsing of brain and behavior may allow us to make greater progress toward understanding the neurobiology of ADHD. A prior *behavioral* observation is that among school-age children, ADHD-associated increases in mirror overflow are particularly evident when volitionally moving their non-dominant hand (MacNeil et al., 2011; McAuliffe et al., 2020). While prior ADHD samples have established greater overflow in the non-dominant hand, there is scant evidence to clarify the physiological basis of this effect. Indeed, there are scant physiological data in ADHD from any modality to clarify how brain control of the non-dominant hand differs from the control of the dominant hand and whether the relationship of dominant to non-dominant hand control differs between groups.

Alpha (8-12 Hz) oscillations in sensorimotor areas are inhibited during movements (Crone et al., 1998). The associated decrease in amplitude are considered to represent the underlying population of neurons for changes in synchronous state (event-related desynchronization [ERD]) (Seeber et al., 2016). We have previously found less asymmetry in alpha ERD in children with ADHD compared to TD controls (McAuliffe et al., 2020). As the alpha rhythm is generally understood to be inhibitory, alpha ERD reflects activation and correlates with metabolic activation of brain regions (Klimesch, 2012; Klimesch et al., 2007; Neuper et al., 2006). These analyses, which combined data from voluntary, unilateral movements in the left and right hands (and mirror overflow in the opposing hand), therefore found atypical *symmetry* of activation in ADHD associated with unilateral hand movement.

The aim of the analysis presented here was to determine whether mirror overflow differed between the dominant and non-dominant hands in ADHD and TD groups and to explore whether and how dominant and non-dominant motor physiology differed between groups. We used data from the same participants as the previous study (McAuliffe et al., 2020). Here, in order to increase our theoretical spatial resolution when assessing the left versus right hemisphere, we used EEG source imaging (He et al., 2019; Luo et al., 2019). Our analysis plan was as follows. We examined (1) group × hand interaction effects of behavioral overflow; (2) group × hand interaction effects of alpha ERD as well as group × hemisphere and group × hand × hemisphere interactions; (3) associations between alpha ERD and behavioral measures of mirror overflow, stratified by dominant and non-dominant hand; and (4) the relationship between ADHD symptom severity and alpha ERD, in order to construct a picture of whether differential control of the non-dominant hand may present a particularly relevant target for further investigation of the neurobiology of ADHD as a whole.

## 2. MATERIALS AND METHODS

The study was conducted in accordance with the World Medical Association code of ethics (Declaration of Helsinki) for experiments involving human subjects. This study was approved by the institutional review board of Johns Hopkins Medicine, Baltimore, MD. Written informed consent and oral assent were obtained from the legal guardians and children, respectively, after the study procedures were fully explained.

### 2.1. Participants and Assessment

The participants were identical to those reported in our previous study (McAuliffe et al., 2020). We recruited 50 children (8-12 years); 25 met the study criteria for ADHD (18 males, mean age=10.36 years, SD=1.24) and 25 were age- and sex-matched typically-developing (TD) controls (19 males, mean age = 10.73 years, SD = 1.33). All participants were right-handed, as evaluated by the Edinburgh Handedness Inventory (Oldfield, 1971).

ADHD diagnosis made using a structured or semi-structured parent interview, either the Kiddie Schedule for Affective Disorders and Schizophrenia (Kaufman et al., 1997) (K-SADS; *n*=38) orthe Diagnostic Interview for Children and Adolescents (Reich, 2000) (DICA-IV; *n*=12); the Conners Parent Rating Scale-Version 3 (Conners, 1997) (*n*=50) and the ADHD Rating Scale (Fabiano et al., 2006) (ADHD-RS; *n*=49) were used to confirm diagnosis and to provide dimensional measures of ADHD symptom severity. Participants were included in the ADHD group if they (1) met criteria for an ADHD diagnosis either on the K-SADS or DICA-IV and (2) received a T-score of 60 or higher on the DSM Inattentive or DSM Hyperactive-Impulsive scales on the Conners Parent or Teacher (when available) rating scale (third edition), or a score of 2 or 3 (i.e., symptoms rated as occurring often or very often) on at least 6/9 items on the Inattentive or Hyperactivity/Impulsivity scales of the ADHD-RS Home or School (when available) Version. ADHD subtype was determined by integrating symptoms endorsed across the diagnostic interview and the Conners and DuPaul’s rating scales. Within the ADHD group, 68% met criteria for the combined subtype and 32% met criteria for the inattentive subtype. Children with ADHD who took stimulant medication (52%) were asked to stop the day before the experiment. Parents were instructed on both the diagnostic interview and report forms to make ratings based on their children’s symptoms off medication. Additionally, children taking psychotropic medications other than stimulant medication did not discontinue their medication for study visits. Two children with ADHD were taking SSRIs at the time of their study visit.

Children with a history of other neurological or psychiatric disorders were excluded. Additionally, those with a full-scale IQ below 80 were excluded in the study, as measured by full-scale IQ (Wechsler, 2014). Children with ADHD showed a trend toward lower IQ than TD controls, as measured by the General Ability Index (GAI) (ADHD mean=112.48±11.41, TD mean=118.72±11.68, *p*=0.062, Cohen’s *d*=0.54). GAI is a composite ability score for the Wechsler Intelligence Scale for Children–fourth edition (WISC-IV) which is used to examine the IQ of children and minimizes the impact of tasks involving processing speed and working memory, known alterations in ADHD (Rowe et al., 2010).

### 2.2. Study Paradigm

TSD131 finger twitch transducers (Biopac Systems Inc., Goleta, CA) were used to quantify mirror overflow in degrees of displacement from a baseline position. Transducers were attached to the metacarpophalangeal joints of the index and ring fingers of the left and right hands to capture finger flexion and extension. The installation of transducers on only these two fingers provides an accurate reflection of overflow of the entire hand, while also increasing the comfort and flexibility of the participants. ACQKnowledge Software 4.2V (Biopac Systems Inc.) was used to calibrate the transducers at 0 °and 45 ° prior to the start of the experiment.

All participants were asked to complete a sequential finger tapping task to assess mirror overflow in the non-tapping hand. Participants were counterbalanced within diagnosis and sex with regard to starting hand (i.e., begin with a left-hand finger-tapping [LHFT] block *vs*. a right-hand finger-tapping [RHFT] block). Five blocks of finger sequencing were collected. Each block consisted of 20 trials (10 LHFT and 10 RHFT) per block. Each trial lasted six seconds, with a one-second rest/baseline period prior to the “Go” cue and five seconds of tapping after the “Go” cue in each trial. Correct positioning showed the tapping hand positioned upright and facing a camera in front of the participant and the non-tapping hand resting over a pillow on the participants’ lap so as not to restrict extension and flexion in their fingers, and thus allowing for overflow movements. Participants were asked to tap their finger pad to the thumb in sequence (one sequence: index-middle-ring-pinky) “as big and fast” as possible to ensure valid, independent taps. Average mirror overflow was computed by averaging the cumulative angular deflection of the non-tapping hand for all trials across blocks together at the same time during LHFT and RHFT. Outlier values (greater than 2 SD) were manually rejected. 94 outliers were removed from 2500 trials total. These outliers affected the TD group less than the ADHD group, but there was no statistical difference between LHFT or RHFT trials.

### 2.3. EEG Acquisition and Analysis

EEG was recorded with an asa-lab amplifier (Advanced Neuro Technologies, Netherlands) and a 47-channel WaveGuard cap system. All electrodes were placed according to the International 10-20 system. Data were recorded at a 1024 Hz sampling rate and down-sampled to 512 Hz. All data were re-referenced to the average of all channels, as per the amplifier design. Impedances were kept below 15 kΩ. EEGLAB toolbox (Delorme and Makeig, 2004) was used for EEG data preprocessing. After the imposition of a 1 Hz high pass finite impulse response (FIR) filter and an 80 Hz FIR lowpass filter, EEG data were segmented into epochs 6 seconds in length, including 1 second before onset (“Go” cue) and 5 seconds subsequently. Baseline correction was performed. A 60 Hz notch filter was used to remove the power interference. We used a blind source separation algorithm to remove artifacts related to eye blink, eye movements and muscle movements(Delorme et al., 2012). Bad channels were interpolated, and bad trials were visually inspected and manually removed. We selected a 1.5-second window for each trial (1.5 to 3 seconds after movement onset) because the mirror overflow movements were most likely to occur during this time period (McAuliffe et al., 2020).

The Brainstorm toolbox (Tadel et al., 2011) was then used for EEG source estimation and ERD analyses. For EEG source imaging analysis, we added EEG positions using the ICBM152 template. To describe the propagation of the electric fields from the cortical surface to the scalp, OpenMEEG was used to formulate the forward model as a boundary element model (BEM) for each participant (Gramfort et al., 2010; Kybic et al., 2005). This symmetric BEM uses three realistic layers, including the scalp, inner skull and outer skull. Electrode positions and BEM were co-registered to match four anatomical landmarks: the vertex, nasion, and left and right preauricular points. Then we computed the noise covariance matrices based on the baseline of individual trials. We used the standardized low-resolution brain electromagnetic tomography (sLORETA) algorithm (Pascual-Marqui, 2002) to solve the inverse problem. After the source imaging analysis, we performed time-frequency decomposition in the EEG source level using the Morlet wavelet algorithm (Morlet et al., 1982). For the Gaussian kernel, the parameters of the mother wavelet were set to a 3-second FWHM with a center frequency of 1 Hz. Finally, ERD (Pfurtscheller and Da Silva, 1999) was calculated. The ERD was calculated using the following formula: 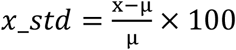, where *x* refers to the data-point to be normalized, and μ refers to the mean over the baseline measurements. We focused on the alpha frequency (8-12 Hz), which was determined by a data-driven approach from the spectrograms. The mu rhythm or sensory-motor rhythm (SMR) is comprised of both alpha and beta components which dissociate in terms of modulation in different experimental conditions (Salmelin et al., 1995). We focused here on the alpha component because of its relevance to altered motor control within this sample in a prior analysis (McAuliffe et al., 2020). The border of the sensorimotor regions was selected via the Broadmann atlas (Maldjian et al., 2003) and included S1, M1, SMA and dorsal premotor cortex. The alpha rhythm was calculated identically across groups and hands.

### 2.4. Statistical Analyses

Statistical analysis was performed using the SPSS Statistics software (version 21.0, IBM Corp., Armonk, NY). We ran independent-samples *t*-tests between ADHD and TD groups to examine potential differences in demographics, behavioral mirror overflow (via goniometer) and clinical data (Conners ADHD Inattention and Conners ADHD H/I). All *t*-tests were 2-sided, with α=0.05. We subsequently adjusted for the potential confound of age, sex and GAI in between-group statistical analyses. All statistical results presented were adjusted for age, sex and GAI.

Because previous datasets have shown an increased burden of mirror overflow movements specifically during voluntary movement of the *non-dominant* hand in ADHD, the primary aim was to show whether previously identified ERD (cortical activation) differences were attributable primarily to tapping of the left hand, within this sample of right-handed participants. Our primary, hypothesis-testing model was a mixed repeated-measures ANOVA; the within-subject factors were “hemisphere” (left, right) and “hand” (LHFT, RHFT) and the between-subject factor “group” (ADHD, TD), with added covariates age, sex and GAI. Additionally, a hand × group interaction effect was of primary importance for hypothesis testing. We further examined group × hemisphere and group × hand × hemisphere interaction effects. We then examined (1) whether there was evidence for a hand (LHFT/RHFT) × group interaction effect in behavioral overflow, as assessed by goniometer; (2) whether the brain (ERD)-behavior (overflow) relationship by hand is similar or different in ADHD as compared with controls, using a moderation analysis (Hayes and Rockwood, 2017); and (3) whether there was a group × hand interaction in the association between ERD and clinical ADHD symptom severity, using Pearson’s *r*.

## 3. RESULTS

### 3.1. Demographics

Table 1 shows the demographics (age, sex and GAI) and clinical examination results (ADHD severity and motor ability scores) of children with ADHD and TD controls.

**Table 1.**
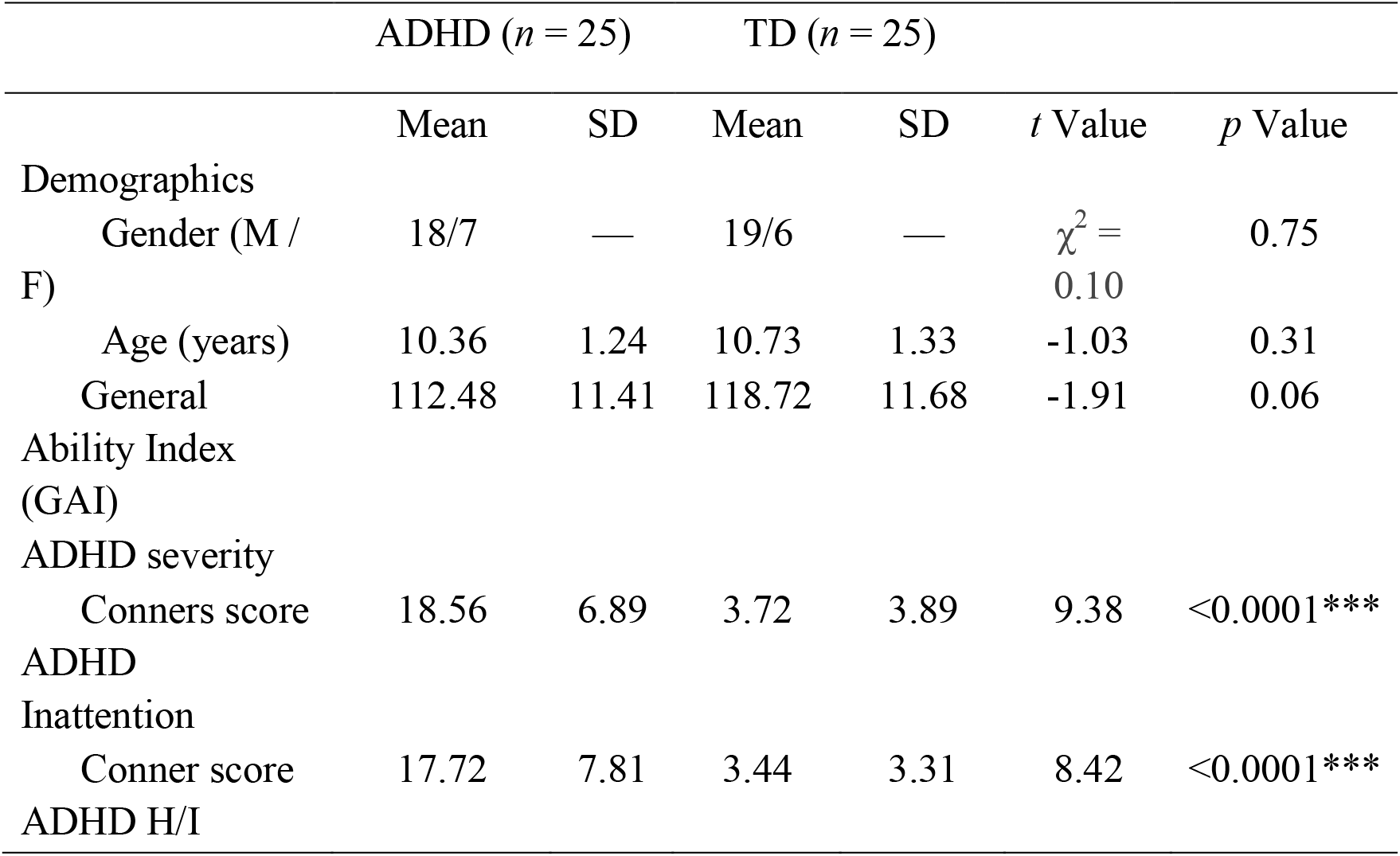
Demographics and behavioral results between the ADHD and TD groups.

### 3.2. Behavioral Overflow

We failed to find a significant group × hand interaction effect for mirror overflow (*p* =0.28, 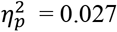), whereas there was a significant main effect of hand (*p* = 0.03, 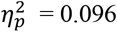) on behavioral overflow, with greater overflow during LHFT (as measured in the right hand). On *post hoc t*-tests, we found that children with ADHD had a significantly greater magnitude of overflow movements compared with TD controls during LHFT (ADHD mean=30.64±16.70, TD mean=17.57±12.75, *t*(48)=3.11, *p*=0.01, Cohen’s *d*=0.88), but the difference during RHFT did not reach statistical significance in this sample (ADHD mean=23.26±14.70, TD mean=15.57 ±18.61, *t*(48)=1.58, *p*=0.25, Cohen’s *d*=0.46) (Figure 1). Further, children with ADHD in this sample showed greater overflow during LHFT than during RHFT (paired *t*-test, *p*=0.046, Cohen’s *d*=0.47) (Figure 1C), whereas no within-group statistical difference in LHFT *vs*. RHFT overflow was observed in TD controls (paired *t*-test, *p*=0.37, Cohen’s d =0.13) (Figure 1D).

**Figure 1.**
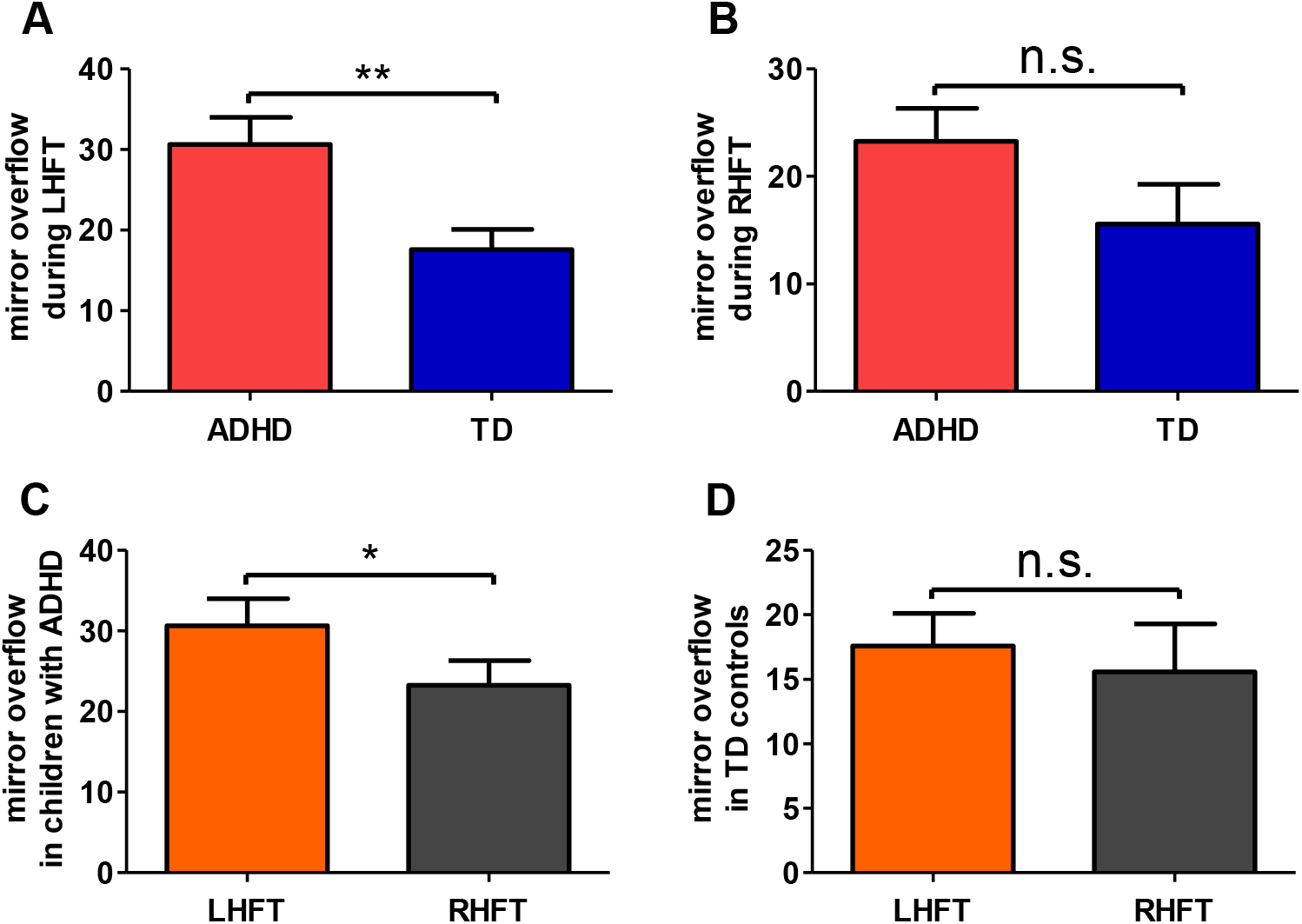
Behavioral mirror overflow results. (A) and (B) showed mirror overflow in children with ADHD and TD controls during LHFT and RHFT, respectively. Children with ADHD demonstrated significantly greater amount of mirror overflow than TD controls *during LHFT only*. (C) and (D) showed mirror overflow during LHFT *vs*. RHFT in both ADHD and TD groups. Children with ADHD had significantly more mirror overflow during LHFT than RHFT, but TD participants did not. \**p*<0.05, \*\**p*<0.01. Error bars are the standard error of the mean (s.e.m.) and n.s. denotes not significant.

### 3.3. Alpha ERD

Note that we use “contralateral” and “ipsilateral” to refer to the cerebral hemisphere *relative to the voluntarily tapping hand*, and *not* relative to the hand producing overflow. Our primary analysis was a mixed repeated-measures ANOVA model, examining for hand (LHFT, RHFT) × group, adjusted for sex, age and GAI. There was a significant group × hand interaction effect in alpha ERD (*p*=0.016, 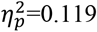). We also looked for but failed to find hemisphere × group interaction effects (*p*=0.82, 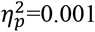) or hand × group × hemisphere interaction effects (*p*=0.40, 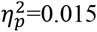). We observed a significant main effect of hand on contralateral sensorimotor areas (*p*=0.031, 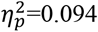).

*Post hoc t-*tests showed significant group differences in alpha ERD during LHFT within both contralateral sensorimotor areas (ADHD mean=-21.93±21.29, TD mean =-35.17±17.05, *t*(48)=2.43, *p*=0.019, Cohen’s *d*=0.69) and ipsilateral sensorimotor areas (ADHD mean=-19.68±17.48, TD mean =-30.08±15.21, *t*(48)=2.24, *p*=0.029, Cohen’s *d*=0.63) (Figures 2 and 3).

**Figure 2.**
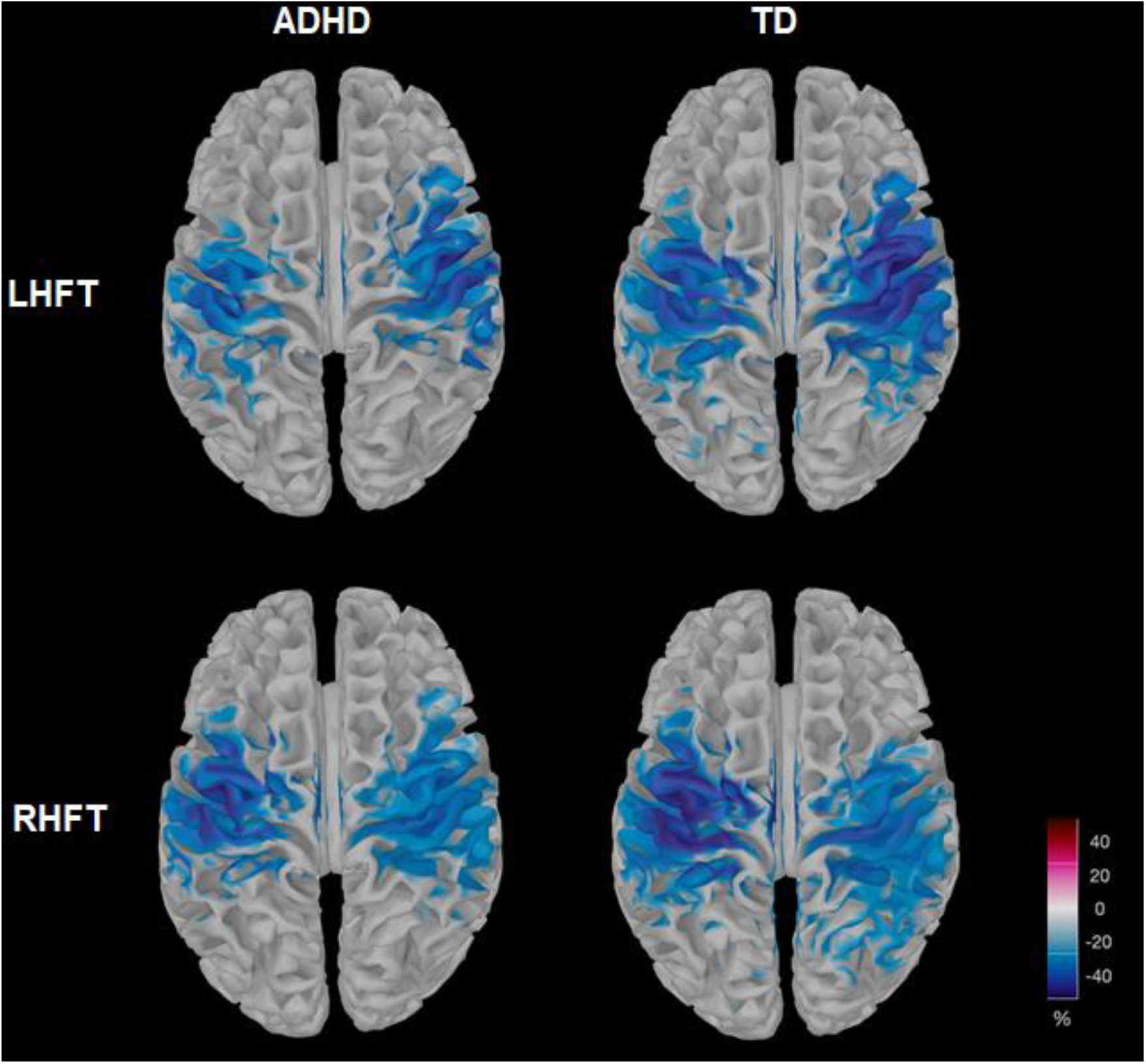
Alpha ERD in children with ADHD and TD controls during LHFT and RHFT. Children with ADHD showed decreased alpha ERD compared with TD controls in contralateral and ipsilateral sensorimotor areas, especially during LHFT.

**Figure 3.**
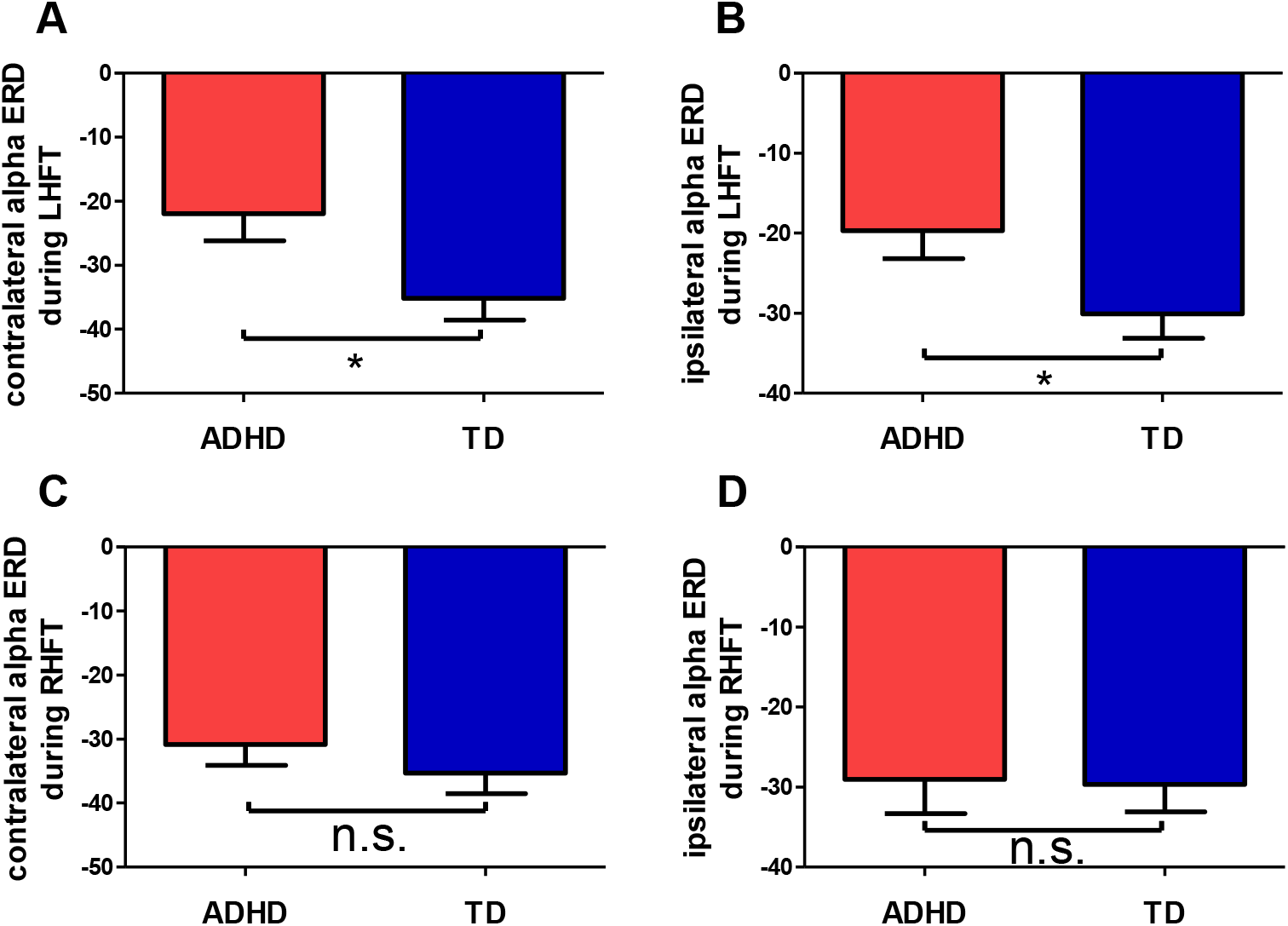
Between-group differences in contralateral and ipsilateral alpha ERD during both LHFT and RHFT in both ADHD and TD group. The two groups differed significantly during LHFT in (A) contralateral alpha ERD in sensorimotor areas, and (B) ipsilateral alpha ERD in sensorimotor areas. No diagnostic effect was observed for RHFT in either (C) contralateral and (D) ipsilateral alpha ERD. In summary, group differences were seen in both ipsilateral and contralateral sensorimotor areas during LHFT but not RHFT. *p<0.05, **p<0.01, and n.s. = not significant. Error bars are the standard error of the mean (s.e.m.).

There was a significant group × hand interaction effect in alpha ERD in ipsilateral sensorimotor areas (*p*=0.022, 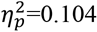). We also observed a significant main effect of hand on ipsilateral alpha ERD (*p*=0.037, 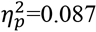). Furthermore, compared to TD controls, children with ADHD demonstrated decreased alpha ERD during LHFT in ipsilateral sensorimotor areas (ADHD mean=-19.68±17.48, TD mean =-30.08±15.21, *t*(48)=2.24, *p*=0.029, Cohen’s *d*=0.63) (Figure 2 and 3). Restated, children with ADHD showed less alpha ERD than TDs in sensorimotor areas, both ipsilateral and contralateral, during LHFT.

In contrast, no statistically significant effect of diagnosis was observed for RHFT in contralateral alpha ERD in sensorimotor areas (*p*=0.33, Cohen’s *d*=0.28) (Figure 3). No significant inter-group differences were found in ipsilateral alpha ERD during RHFT (*p* = 0.91, Cohen’s *d*=0.03)(Figure 3).

Within the ADHD group, there were significant differences between LHFT and RHFT in contralateral alpha ERD in sensorimotor areas (*p*=0.01, Cohen’s *d*=0.47)(Figure 4). Children with ADHD showed significantly lower alpha ERD in the contralateral sensorimotor areas during LHFT than RHFT. Furthermore, there were also significant differences within the ADHD group between LHFT *vs*. RHFT in ipsilateral alpha ERD in sensorimotor areas (*p*=0.005, Cohen’s *d*=0.47) (Figure 4).

**Figure 4.**
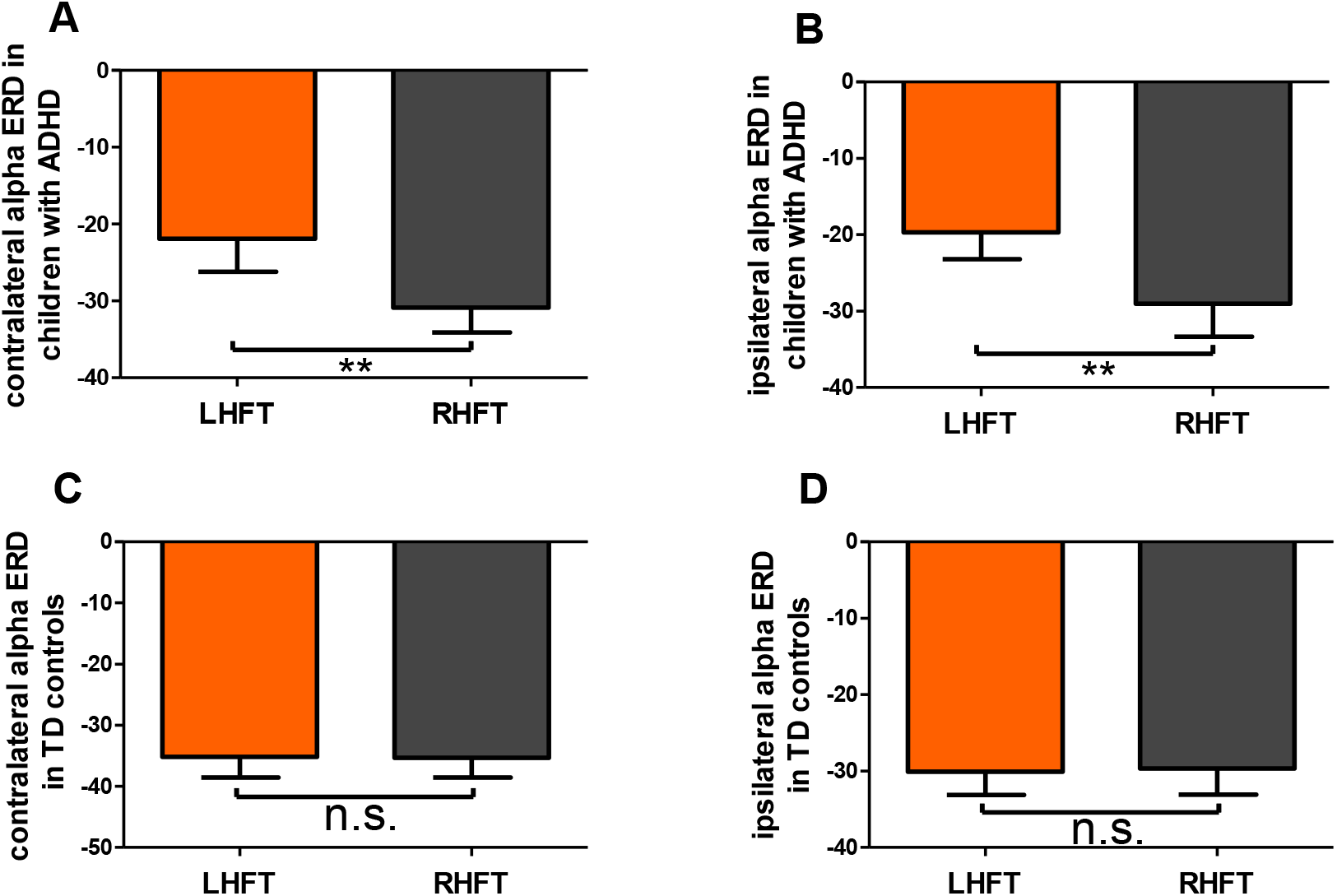
Within-group, between-hand differences in contralateral and ipsilateral alpha ERD in sensorimotor areas. Children with ADHD showed significant differences in contralateral (A) and ipsilateral alpha ERD (B) in sensorimotor areas between LHFT *vs*. RHFT, whereas TD controls did not show significant differences in contralateral (C) or ipsilateral alpha ERD (D) between LHFT *vs*. RHFT. *p<0.05, **p<0.01, and n.s. = not significant. Error bars are the standard error of the mean (s.e.m.).

And the ipsilateral ERD during LHFT was lower than ERD during RHFT. Within the TD group, no significant differences were found between LHFT *vs*. RHFT in contralateral (*p* = 0.95, Cohen’s *d* =0.01) or ipsilateral alpha ERD (*p* =0.88, Cohen’s *d* =0.03) (Figure 4).

In conclusion, children with ADHD demonstrated significant differences compared with TDs in both ipsilateral and contralateral ERD during LHFT but not RHFT (Figure 3). For children with ADHD, alpha ERD was significantly lower for non-dominant (LHFT) than for dominant (RHFT) finger tapping. By contrast, TD controls showed no statistical difference by dominant *vs*. non-dominant hand (Figure 4). Significant between-group differences were seen only during LHFT.

### 3.4. Relationship Between Overflow and ERD

Having found group differences in overflow only in the dominant hand (right) during non-dominant hand movements (LHFT), and having found group differences in ERD only during LHFT, we next examined the statistical relationship between ERD and overflow. Behavioral overflow was significantly correlated with contralateral alpha ERD in sensorimotor areas during LHFT within the ADHD group (*r*=-0.60, *p=*0.004), whereas no significant correlation was found during RHFT (*r*=0.02, *p=*0.93) (Figure 5). Moreover, there were no significant correlations between overflow and ipsilateral alpha ERD in sensorimotor areas during LHFT or RHFT within the ADHD group (LHFT: *r*=-0.27, *p*=0.23; RHFT: *r*=0.15, *p*=0.54) (Figure 5). Furthermore, there were no significant correlations between overflow and contralateral or ipsilateral alpha ERD in the TD group during LHFT and RHFT (Figure 5). Noting that the regression line appeared to have a different slope between LHFT and RHFT (Figure 5). A moderation analysis was performed to investigate the moderation effect of hand on ERD-overflow relationship. We identified a statistically significant moderation effect of hand within the contralateral alpha ERD in ADHD (*p*=0.04), whereas there was no moderation effect of hand within the ipsilateral alpha ERD in ADHD (*p*=0.36). Restated, in the ADHD group, the relationship between overflow and ERD is different when the individual is tapping the left hand *vs*. the right hand.

**Figure 5.**
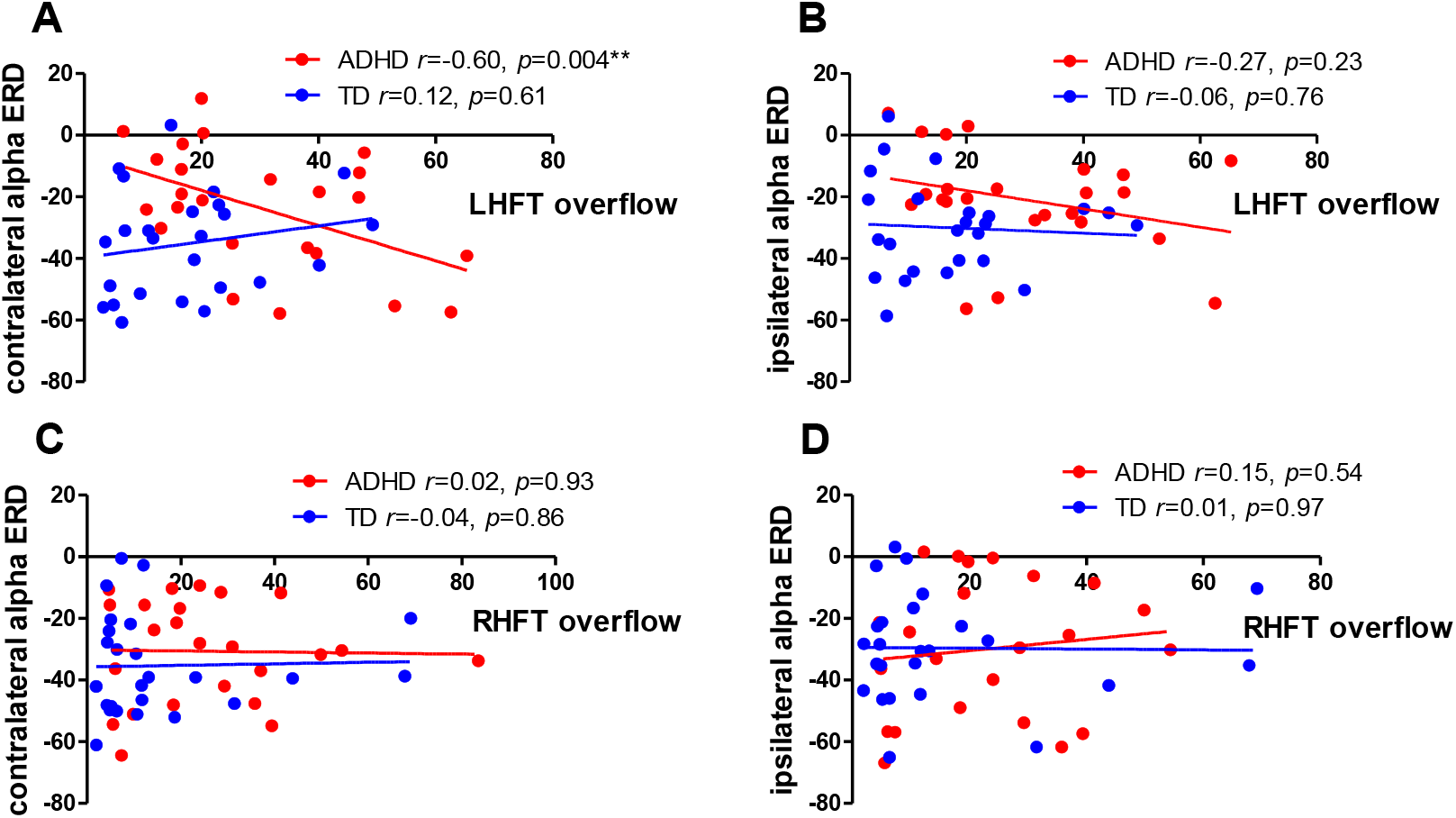
Correlations between contralateral alpha ERD and behavioral overflow during LHFT and RHFT in children with ADHD *vs*. TD controls. (A) and (B) showed correlations between behavioral overflow and contralateral or ipsilateral alpha ERD during LHFT in the ADHD and TD group. (C) and (D) showed correlations between behavioral overflow and contralateral or ipsilateral alpha ERD during RHFT in the ADHD and TD group. Note that the regression lines have different slopes by group for LHFT but not for RHFT; restated, the relationship between overflow (behavior) and contralateral ERD (brain) is different for children with ADHD *vs*. TD controls, but only for LHFT (moderation effect *p*=0.04). Significant correlation was seen in the relationship between overflow and contralateral ERD in sensorimotor areas (*p* = 0.004).

No moderation effect was seen in TD controls (contralateral ERD: *p* =0.50; ipsilateral ERD: *p*=0.90). These statistical results were adjusted for age, sex and GAI. In summary, TD controls did not show astatistical association between alpha ERD and overflow. Children with ADHD showed an association between ERD and overflow, but only during LHFT. More specifically, the only association was between mirror overflow in the right hand (during volitional movement of the left hand) and ERD in the right hemisphere (specifically in the sensorimotor areas; more ERD = more overflow).

### 3.5. Association Between ERD and ADHD Clinical Severity

Figure 6 demonstrates the relationship between ADHD symptoms and alpha ERD. Among children with ADHD, there was significant correlation between Conners ADHD Hyperactivity/Impulsiveness (H/I) and LHFT contralateral alpha ERD in sensorimotor areas (*r*=0.43, *p*=0.04). There were no correlations between ipsilateral alpha ERD and Connors ADHD scales in the ADHD group during LHFT. Moreover, there was no correlation between either Connors ADHD subscale score and ERD in the sensorimotor areas of either hemisphere during RHFT in ADHD. There were no significant correlations with either Connors ADHD subscale score and contralateral or ipsilateral ERD in sensorimotor areas, during either RHFT or LHFT within the TD group. In summary, not only was an atypical physiology-overflow relationship seen during LHFT in the ADHD group—a relationship that was different than the one seen in ADHD-RHFT and TD for either hand—but also, contralateral ERD during LHFT in the ADHD group is the only physiological measurement that correlated with ADHD symptom severity.

**Figure 6.**
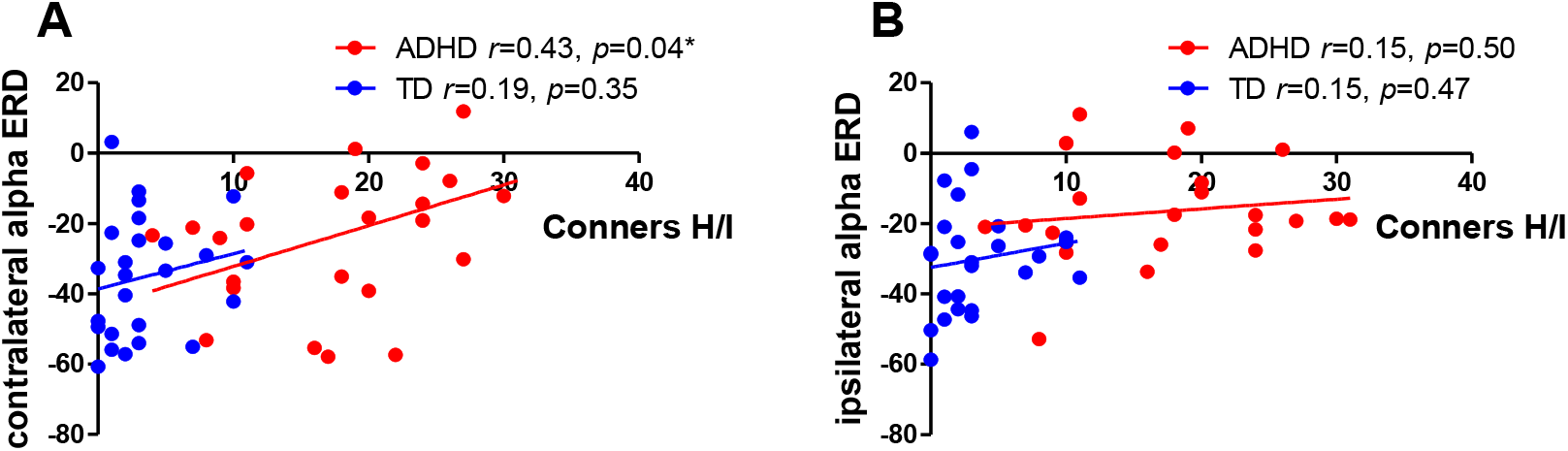
Correlations between ADHD symptoms and alpha ERD. (A) showed correlations between contralateral alpha ERD during LHFT and Conners ADHD Hyperactivity/Impulsiveness (Conners H/I) in the ADHD and TD group. (B) showed correlations between ipsilateral alpha ERD during LHFT and Conners H/I in the two groups. There were significant correlation between contralateral alpha ERD and Conners H/I in children with ADHD during LHFT, whereas the contralateral alpha ERD of TD controls did not show a linear relationship with Conners ADHD Hyperactivity/Impulsiveness.

## 4. DISCUSSION

Our primary finding was that previously described, ADHD-related differences in alpha ERD during finger tapping (McAuliffe et al., 2020) are driven by the effect of the voluntary movement of the non-dominant hand. Moreover, group differences in overflow are also driven by the overflow that occurs during voluntary tapping of the left hand in the right-handers (per *post hoc* statistical tests). That mirror overflow is increased during voluntary movement of the non-dominant hand has been reported in other clinical conditions (Armatas et al., 1994, 1996; Hoy et al., 2004) (but also see (Cernacek, 1961)). While the group × hand differences in ERD were seen in both hemispheres, we note that it was the hemisphere contralateral to the tapping hand (ipsilateral to the hand demonstrating mirror overflow) that showed a statistical association with magnitude of overflow and with H/I symptoms.

What do these results tell us about the generation of overflow movements in ADHD? We can consider our results using a framework of alternative hypotheses developed in the broader mirror overflow literature (beyond only ADHD), which has identified two primary hypotheses to explain how mirror movements are generated (Hoy et al., 2004) (Figure 7). The Transcallosal Hypothesis states that mirror overflow during LHFT would occur as follows: activation of the motor system in the right hemisphere (“contralateral” to volitional movement, in the labeling system we have used here) simultaneously generates both motor commands that run down the decussating corticospinal tract (CST) to the left hand as well as inhibitory signals that pass through the corpus callosum to the left hemisphere (“ipsilateral”), suppressing potential mirror movements in the right hand. When mirror overflow occurs in the right hand, it is caused by ineffective transcallosal inhibitory signals from the contralateral to ipsilateral motor cortex (Addamo et al., 2007; Beaulé et al., 2012; Cox et al., 2012; Gaddis et al., 2015). Indeed, there is considerable prior data to suggest ADHD is associated with alterations of interhemispheric inhibition (Mostofsky et al., 2003; Mostofsky et al., 2006; Wu et al., 2012). However, if the Transcallosal Hypothesis were true, we would have expected in this dataset to see a strong statistical relationship between the degree of activation (here: alpha ERD) in the left hemisphere (“ipsilateral” to primary movements) and the magnitude of mirror movements in the right hand during LHFT, since it is decreased inhibition of the left hemisphere motor system that generates decussating motor commands to the right hand. This pattern of results was not seen.

**Figure 7.**
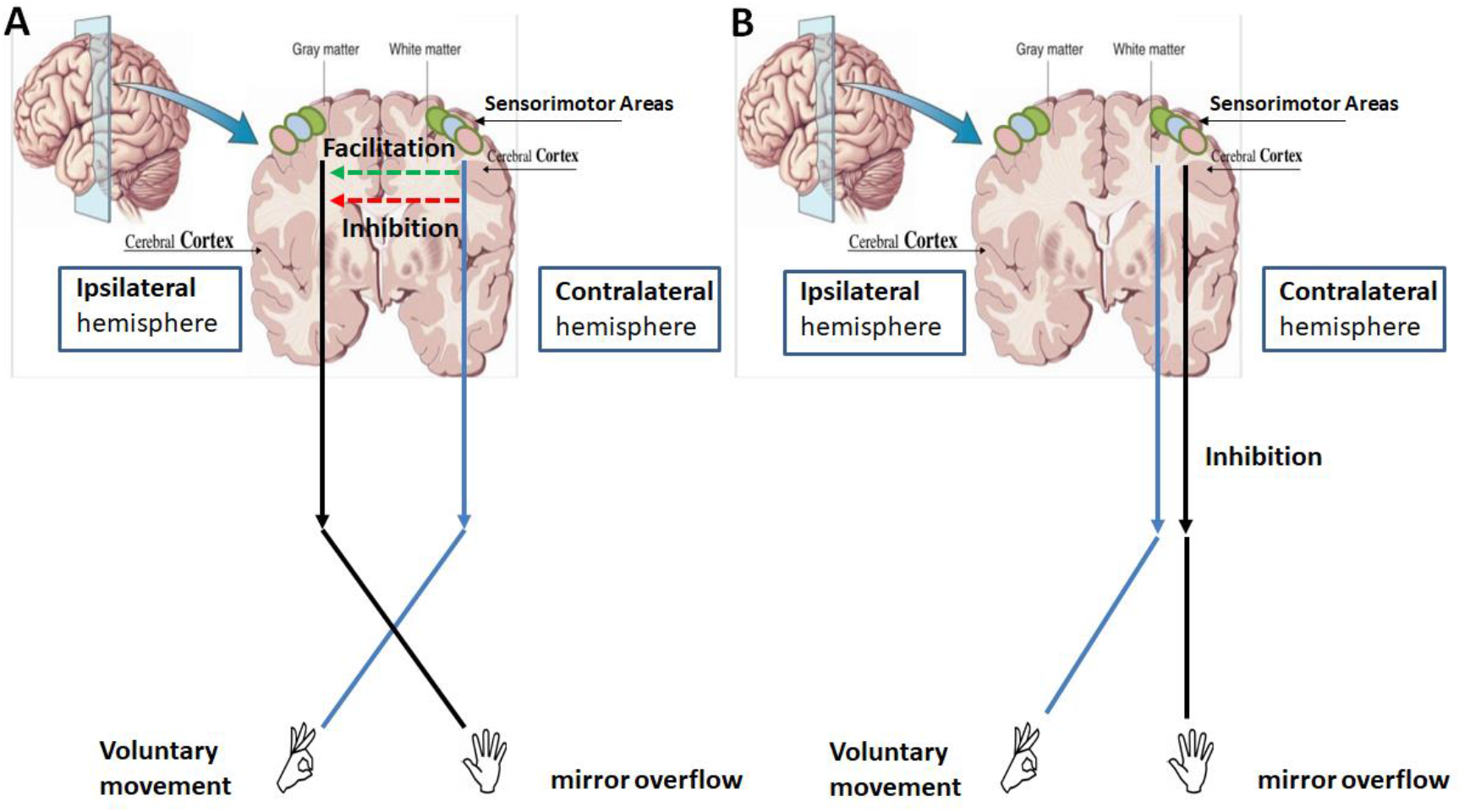
Mirror overflow models. (A) showed the Transcallosal Hypothesis, (B) showed the Ipsilateral CST Hypothesis. Contralateral hemisphere and ipsilateral hemisphere in the figure refer to contralateral/ipsilateral to the voluntary movement, whereas “ipsilateral” in the title of the Ipsilateral CST Hypothesis refers to non-decussating CST fibers that travel to the hand with overflow from the hemisphere ipsilateral to the overflow hand.

An alternative account in the mirror overflow literature is the Ipsilateral (i.e., non-decussating) CST Hypothesis, which states that, while decussating fibers carry motor signals from the right hemisphere to the left hand, non-decussating CST fibers carry motor signals from the right hemisphere to the right hand. In this case, greater activation of the right hemispheric motor system (greater alpha ERD) would be associated with greater mirror overflow. And the inhibition is weaker during the non-dominant hand movements. Consistent with predictions of this hypothesis, contralateral ERD was in fact correlated with behavioral overflow. This result, deciding between the Transcallosal Hypothesis and the Ipsilateral CST Hypothesis, is a mores specific question than we asked in our prior analysis of these data (McAuliffe et al., 2020).

Why might it be that voluntary movement of the *non-dominant* hand produces greater mirror overflow in ADHD, but voluntary movement of the dominant hand does not? One possibility is that voluntary movements of the non-dominant hand require more attention to be performed successfully. Increased attention directed toward the non-dominant, voluntarily tapping hand (left) somehow takes away from the inhibition on the cortical regions that generate mirror overflow in the right hand. However, in the absence of direct measurement of this possibility (e.g., performing finger-tapping during an interfering task), if this hypothesis were true, we may have expected a correlation between severity of ADHD symptoms and degree of overflow within the ADHD group. We did not find such a correlation.

Another account suggests differential excitability of the dominant *vs*. non-dominant cortical regions, driven by greater transcallosal inhibition of the non-dominant hemisphere by the dominant, rather than *vice versa* (Beaulé et al., 2012). Transcranial magnetic stimulation (TMS) studies of right-handers have reported that the modulation of interhemispheric inhibition was asymmetrical during the dominant *vs*. nondominant hand movements (Duque et al., 2007). For example, for the right-handed participants, the inhibition effect is greater from the dominant M1 (in the left hemisphere) to the non-dominant M1 (in right hemisphere) than vice versa (Beaulé et al., 2012; Netz et al., 1995; Serrien et al., 2006; Todor and Lazarus, 1986). In short, greater activation of the non-dominant hemisphere is needed to generate similar quality voluntary movements than compared with the activation of the dominant hemisphere to generate dominant-hand movements. This “excess” activation leads to overflow. We are limited in evaluating this hypothesis, first and foremost because this account endorses mirror overflow being driven by transcallosal inhibition, which the current data do not support.

Furthermore, previous studies support a dominant role for the left hemisphere in the control of both right and left hand movements, whereas the right hemisphere controls left-hand movements in right-handers (Serrien et al., 2006). It may be that the left hemisphere is dominant for motor skills in either hand (Serrien et al., 2006), therefore the RHFT and the overflow movements in the left hand can be well controlled by the left hemisphere. On the contrary, because the right hemisphere only controls the movements of left hand, we observed excessive mirror overflow in the right hand. Future studies investigating both left- and right-handed individuals would help to better disambiguate the effect of dominant/non-dominant hemisphere from right/left hemisphere.

In summary, our results focus the study of mirror overflow in ADHD toward the study of overflow occurring during volitional movement of the non-dominant hand. Both behavioral measures of overflow and brain physiology (alpha ERD) show diagnostic-group-related differences during tapping of the non-dominant hand only. These analyses advance beyond our previous analyses by demonstrating that increased ERD (activation) only in the hemisphere ipsilateral to the overflow movements (i.e., contralateral to the volitional movement) result in a greater magnitude of overflow movements, consistent with the Ipsilateral (non-decussating) CST Hypothesis of mirror overflow genesis. Just as the right hemisphere motor ROI ERD correlates with overflow in the ADHD group, it also correlates with H/I symptom severity in the same group. We do not currently have a framework for understanding why right-hemisphere motor region ERD would associate with H/I symptom severity.

Although our results are inconsistent with an Altered Transcallosal Inhibition account of mirror overflow generation in ADHD, we recognize that the literature holds substantial evidence for alterations of transcallosal inhibition in ADHD (Gaddis et al., 2015; Mostofsky et al., 2006). However, we note that Mostofsky *et al*. (2006) found group differences in fMRI activation of the contralateral M1 to the volitionally tapping hand (and ipsilateral to overflow). While Gaddis et al. (2015) found bilateral between-group differences in fMRI activation, it was the fMRI activation in the *contralateral* hemisphere that correlated with overflow. The results of both studies were similar to ours in this regard.

Multi-modal research is needed to reconcile our results, in support of the Ipsilateral CST Hypothesis, with existing evidence in the literature for transcallosal atypicalities in ADHD, as well as to explore why the right hemisphere shows both atypical activation and strong associations with overflow and H/I symptoms. Such research within the same sample of participants may include, in addition to ERD recording, TMS measures of cortical excitability in both hemispheres and anatomical and TMS direct measures of inter-hemispheric inhibition. Additionally, measurement of performance parameters in the voluntarily tapping hand can be modeled in to understand the effects of effort (relevant to some accounts of dominant/non-dominant performance) and the relationships between physiology and voluntary movement performance.

Finally, machine learning based individualized prediction method could be leveraged in the future study to predict the severity of ADHD symptoms in unseen individuals using the alpha ERD data (Cui et al., 2020).

## Acknowledgements

The authors gratefully acknowledge Jaime Lush for his contribution to the acquisition of data. This work was supported by the National Institutes of Mental Health at the National Institutes of Health (R01 MH078160-08S1 and R01 MH085328) to SHM.

